# Image velocimetry and spectral analysis enable quantitative characterization of larval zebrafish gut motility

**DOI:** 10.1101/169979

**Authors:** Julia Ganz, Ryan P. Baker, M. Kristina Hamilton, Ellie Melancon, Parham Diba, Judith S. Eisen, Raghuveer Parthasarathy

**Author notes:** These authors contributed equally (Julia Ganz, Ryan P. Baker). Corresponding authors (JSE,; RP,).

## Abstract

**Summary Statement:** We present a new image analysis technique using image velocimetry and spectral analysis that returns quantitative measures of gut contraction strength, frequency, and wave speed that can be used to study gut motility and other cellular movements.

**Abstract:** Normal gut function requires rhythmic and coordinated movements that are affected by developmental processes, physical and chemical stimuli, and many debilitating diseases. The imaging and characterization of gut motility, especially regarding periodic, propagative contractions driving material transport, are therefore critical goals. Whereas previous image analysis approaches have successfully extracted properties related to temporal frequency of motility modes, robust measures of contraction magnitude remain elusive. We developed a new image analysis method based on image velocimetry and spectral analysis that reveals temporal characteristics such as frequency and wave propagation speed, while also providing quantitative measures of the amplitude of gut motions. We validate this approach using several challenges to larval zebrafish, imaged with differential interference contrast microscopy. Both acetylcholine exposure and feeding increase frequency and amplitude of motility. Larvae lacking enteric nervous system gut innervation show the same average motility frequency, but reduced and less variable amplitude compared to wild-types. Our image analysis approach enables insights into gut dynamics in a wide variety of developmental and physiological contexts and can also be extended to analyze other types of cell movements.

## Introduction

Proper gut motility is vital for the health of many organisms, yet measurement and characterization of motility patterns remains challenging, a consequence of both the diversity of gut phenotypes and the limitations of existing analysis and imaging methods. A variety of disorders can alter dynamics of the gut. In humans, for example, inflammatory bowel disease, irritable bowel syndrome, chronic intestinal pseudo-obstruction, Hirschsprung disease, and other ailments typically cause gut dysmotility (Brosens et al., 2016; Goldstein and Nagy, 2008; Heanue and Pachnis, 2007). Even within a healthy individual, the gut exhibits different types of movements depending, for example, on its digestive state (Furness, 2006; Huizinga and Lammers, 2009). When fasting, the gut experiences the cyclic sweeping patterns of the migrating motor complex (MMC) (Deloose et al., 2012; Furness, 2006; Olsson and Holmgren, 2011). The presence of food triggers changes in gut movements that in turn affect the ingested material. Standing contractions serve to mix and break up food, whereas propagating contractions transport contents along the gut (Furness, 2006; Olsson and Holmgren, 2011; Wood, 2008).

Rhythmic smooth muscle contractions are orchestrated by an interplay between the slow waves of pacemaker-like interstitial cells of Cajal and the enteric nervous system (ENS) (Furness, 2006; Olsson and Holmgren, 2011; Sanders et al., 2014). Gut movements arise from coordinated activation of sensory neurons, as well as both inhibitory and excitatory motor neurons that can be activated by mechanical or chemical stimuli, guided also by gut-extrinsic innervation (Furness, 2006; Olsson and Holmgren, 2011). Although the neuronal circuits and neuronal subtypes that locally regulate contractions have been identified in mammalian models (Furness, 2006; Wood, 2008), little is known about how these different neuronal subtypes work together to coordinate and switch between all of the complex motions of the gut and how gut motility is influenced at the whole organ level by digestive states or other chemical or physiological perturbations (Furness et al., 2014; Wood, 2008).

Our ignorance stems in part from a challenge inherent to the study of gut motility: the gut displays a large variety of dynamic behaviors, yet understanding these behaviors calls for simple and comprehensible characterizations of their parameters. A common analysis method involves the generation of spatiotemporal maps (STMaps) from video data (Hennig et al., 1999; Janssen, 2013). In an STMap, intensity in a two-dimensional (2D) video series is averaged over the short dimension of the gut, giving a one-dimensional measure that varies over time. This is convenient to plot, as the one spatial and one temporal dimension are readily assembled into a two-dimensional graph. Correlated patterns, such as traveling waves along the gut, appear as streaks in the plot. An STMap enables straightforward determination of three important parameters of gut motility: the peristaltic frequency, the propagation velocity for peristaltic waves traveling along the gut, and the wavelength of contractions (Hennig et al., 1999; Holmberg et al., 2004; Holmberg, 2003; Janssen, 2013). A major limitation of STMaps, however, is that they provide at best only qualitative measures of contraction strength, since the image intensity axis is just a measure of brightness, not a quantitative measure of gut shape or motion. In other words, differences in intensity in an STMap cannot be mapped onto measures of the actual magnitudes of gut tissue displacement. The magnitude of contractions is likely to be modulated during both normal gut function and various disease states, thus good measures of this characteristic are needed. More generally, expansion of the repertoire of parameters beyond a basic set of three would allow finer characterizations of different physiological states that are beyond the reach of current methodologies.

In this study, we report a new image analysis technique that returns quantitative measures of gut contraction strength, as well as frequency and wave speed. This approach, described in more detail below, involves applying well-established image velocimetry techniques to videos of gut motility, and analyzing the magnitude of dominant periodic modes via Fourier transformation. The code is freely available on github: https://github.com/rplab/Ganz-Baker-Image-Velocimetry-Analysis. All data values plotted in figures 2-4 and supplementary figure 1-2 are tabulated in the supplemental text file of comma-separated values “supplemental data 1”.

We apply and assess this technique using images of larval zebrafish guts obtained from a custom-built differential interference contrast microscope (DICM) (Baker et al., 2015). Zebrafish are ideally suited for *in vivo* imaging due to their external development and optical clarity during embryonic and larval stage. In addition, zebrafish is an important animal model for studying gut development and function, including aspects of human gut diseases (Ganz et al., 2016; Zhao and Pack, 2017) and gut microbiota function and dynamics (Ganz et al., 2016; Rolig et al., 2017; Wiles et al., 2016). DICM provides high contrast and high resolution optical sectioning, therefore enabling robust image velocimetry calculations. Our method can be more generally used, however, and should, for example, be applicable to dissected preparations commonly used in studies of mammalian guts. Also, as our method is agnostic as to which type of images are analyzed, it can be used for a variety of cellular movements.

To validate our methods, we examine the effects on larval zebrafish gut motility parameters of a chemical stimulus, a physical perturbation, and a biological deficiency, namely acetylcholine, food, and absence of an enteric nervous system, respectively. We find that acetylcholine-treated larvae show a previously reported increase in contraction frequency (Holmberg et al., 2004; Shi et al., 2014) as well as a newly reported increase in contraction amplitude. Comparing gut motility parameters in fed versus unfed larvae, we find that feeding increases contraction frequency and sustains higher amplitudes over the observed developmental window. Zebrafish larvae lacking ENS innervation show decreased contraction amplitude and also reduced parameter variability compared to wild-type siblings. In addition, imaging over longer intervals reveals highly variable gut motility patterns within individual zebrafish larvae that appear to be ENS-dependent, as the variability of these patterns is lower in mutants lacking ENS innervation. We suggest that our analysis method opens exciting new avenues for studying gut motility in zebrafish and other systems.

## Results

### An image analysis technique based on quantitative spatiotemporal maps and spectral analysis identifies gut motility parameters

To distill complex images of gut motility into concise yet meaningful parameters, we developed a new image analysis approach using image velocimetry and spectral analysis (Fig. 1). A typical zebrafish imaged at 6 days post fertilization (dpf) is shown in Fig. 1A. A full description of the technique can be found in the Materials and Methods section; we provide a summary here. In our experiments, videos of zebrafish gut motility were obtained with DICM (Fig. 1B, left column). A velocity map of the material in each image in the series was determined by digital Particle Image Velocimetry (PIV) (Willert and Gharib, 1991). We used well-established and freely available PIV code (Thielicke, 2014) that divided each image into a grid of sub-images; the sub-image pairs in adjacent frames that were maximally correlated with each other revealed the frame-to-frame displacement of material in that region, or equivalently its velocity (Fig. 1B, middle column). Areas outside the gut were discarded from the analysis.

**Figure 1:**
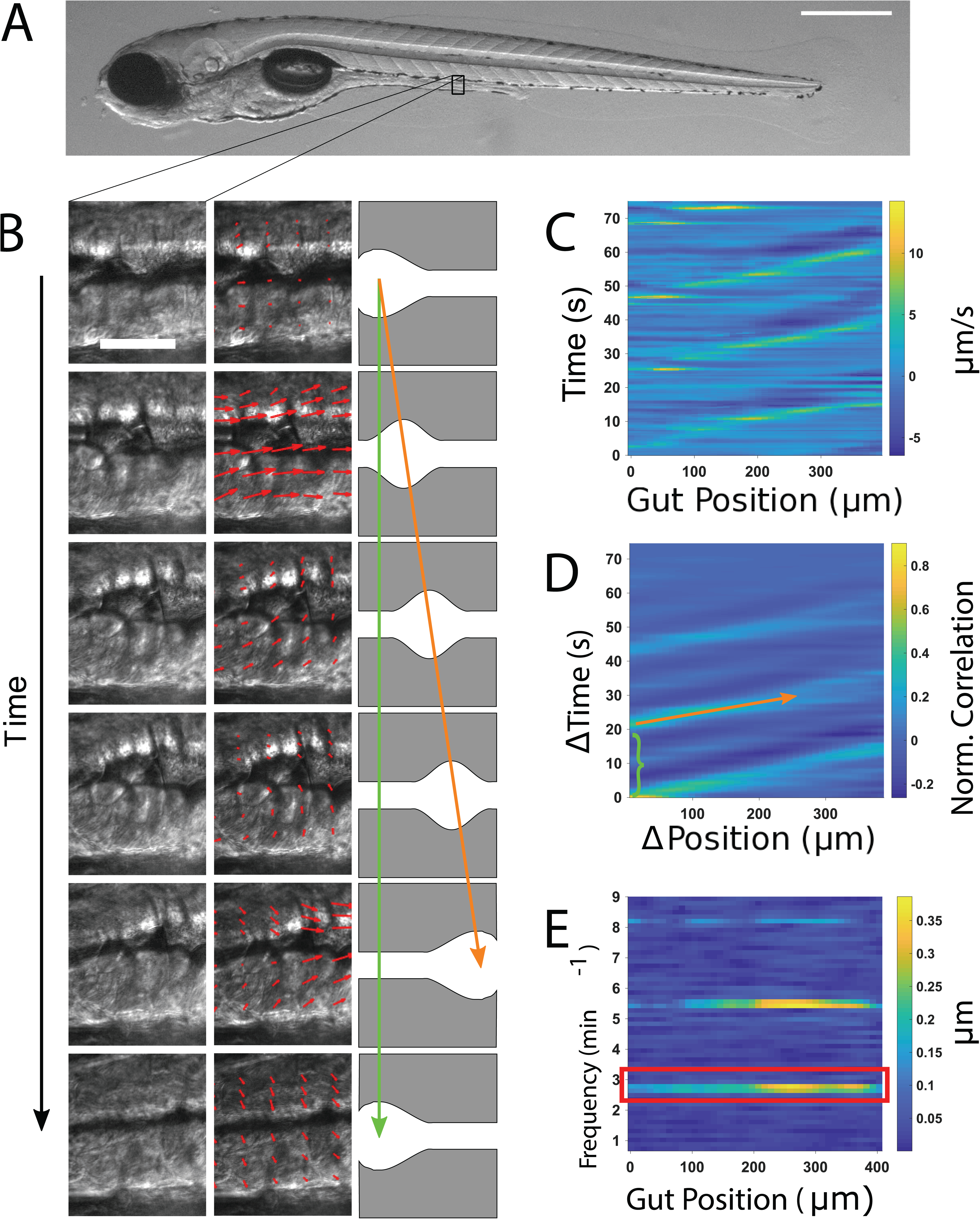
Illustration of gut motility analysis. (A) Brightfield image of a 6 dpf zebrafish larva. Scale bar: 500μm. (B) Left column: a representative series of DIC images of a small region of the midgut. Scale bar 25 μm. Center column: the velocity vector field (red arrows) obtained by performing Particle Image Velocimetry (PIV) on the image series. Right column: schematic illustration of coordinated movements from left to right (anterior to posterior). Wave speed is indicated by the slope of the orange arrow and periodicity indicated by the extent of the green arrow. (C) Quantitative Spatiotemporal Maps (QSTMaps) are obtained by averaging the anterior-posterior component of the velocity vector field along the dorsal-ventral direction and then plotting the curve over time. The color axis represents the velocities, with positive and negative values denoting posterior and anterior movement, respectively. (D) An averaged cross correlation of the QSTMap more clearly reveals motility parameters. The period of motility events corresponds to the time of the first local maximum at ΔPosition = 0 autocorrelation (green bracket; analogous to the length of the green arrow in Fig. 1B). The wave speed corresponds to the inverse slope of maxima in the plot (orange arrow; analogous to the orange arrow in Fig. 1B). (E) A power spectrum of the QSTMap shows the magnitude of gut motion at various frequencies. The amplitude of motility events is determined from the average of the power spectrum at the motility frequency (red box).

Because we were primarily concerned with motion along the anterior-posterior (AP) axis, and its variation along that axis, we considered only the AP components of the resulting two-dimensional displacement map, and further condensed these by averaging along the dorsal-ventral axis (DV). We thereby obtained a one-dimensional curve representing the instantaneous AP frame-to-frame displacement of gut tissue as a function of the distance along the gut. Evaluating this over time, we generated a quantitative spatiotemporal map (QSTMap) of AP displacement as a function of AP position and time (Fig. 1C).

The QSTMap has similarities to STMaps used in previous studies [e.g. (Hennig et al., 1999; Holmberg et al., 2004; Holmberg, 2003; Janssen, 2013)]. The frequency of gut motility events can be inferred from their temporal spacing (Fig. 1B, green arrow, and 1D, green bracket), and the wave speed is given by the slope of linear features in the map (Fig. 1B, orange arrow, and 1D, orange arrow). Unlike STMaps, the intensity of a QSTMap at any point is not simply a measure of image intensity, but rather gives the instantaneous velocity, related to the amplitude of motility events, which we make use of below.

To more robustly quantify wave frequency and speed, we calculated the cross-correlation of the QSTMap: at each AP position (*x*) and time (*t*), we calculated the product of the QSTMap value and its value at a position and time shifted by (Δ*x*, Δ*t*), and then average over all *x* and *t* (Fig. 1D). A wavelike mode of velocity *v*, for example, will be well-correlated with an image of itself shifted by Δ*x* = *v* Δ*t*, while random motions will, on average, be uncorrelated. The time shift of the first local maximum at Δ*x* = 0 represents the periodicity of gut motility (green bracket in Fig. 1D). The inverse slope of the peaks in the cross-correlation map corresponds to the wave speed (orange arrow in Fig. 1D). Parameters such as the wave duration and the variance of wave speed could also be determined.

To characterize the amplitude of gut motility events, not possible with standard methods, we applied spectral analysis to the QSTMap, highlighting periodic signals and quantifying their magnitude. We calculated the one-dimensional Fourier transform of the QSTMap displacement at each AP position (*x*), decomposing the time-varying function into contributions from each of the range of possible frequencies. The square of the Fourier transform, known as the power spectral density, is composed of strong peaks at the frequencies of gut motility events (Fig. 1E), namely the primary frequency (red box) and its harmonics. We defined the gut motility amplitude as the average of the magnitude of the Fourier transform at the primary event frequency.

### Acetylcholine increases the frequency and amplitude of gut motility in zebrafish larvae

The neurotransmitter acetylcholine (ACh) has been shown to increase the frequency of movements in the developing zebrafish gut at several different developmental stages (Holmberg et al., 2004; Shi et al., 2014). To test our image analysis method in an experimental setting with an expected outcome, we treated 6 dpf wild-type larvae with 2.5 mg/ml ACh and compared their gut motility with that of untreated siblings. DICM videos were taken at 5 frames per second (fps) for 5 minute durations and analyzed as described above. In agreement with Shi and colleagues (2014), frequencies were generally higher for ACh-treated larvae than for controls (Fig. 2A), with mean ± s.e.m. values 2.38 ± 0.03 min^-1^ and 2.23 ± 0.05 min^-1^, respectively. In particular, only a few ACh-treated larvae showed frequencies that were lower than the median of the frequencies for untreated, control larvae, and the standard deviation of the motility frequencies for ACh-treated larvae was also lower than for the control larvae (Fig. 2A). The ratio of the mean frequencies for treated and untreated larvae in our experiments is 1.07 ± 0.03, clearly greater than 1, as was also the case in Shi *et al.*’s study in which frequency was assessed from manual counting of occurrences of folds in the gut (Shi et al., 2014)

**Figure 2:**
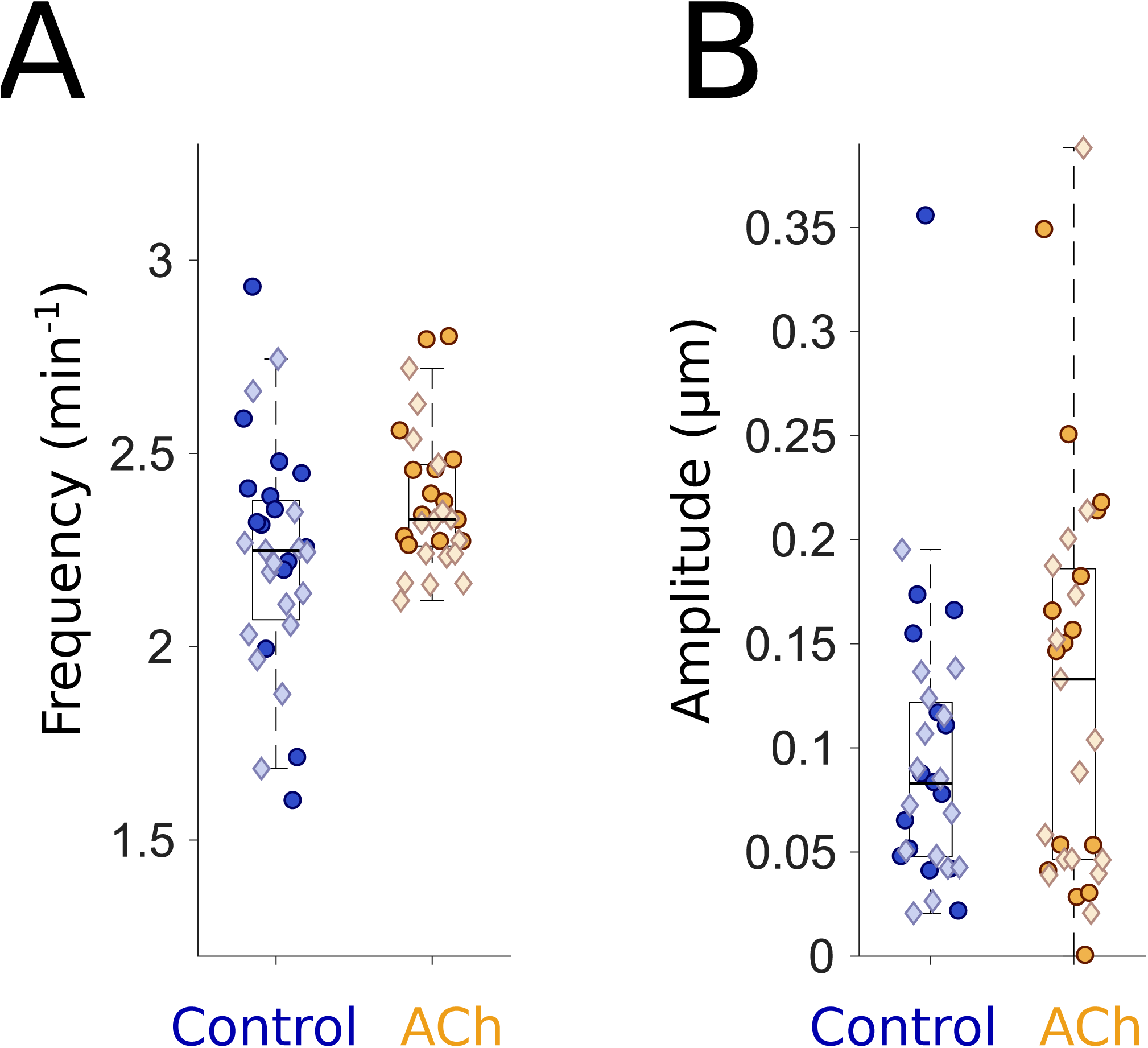
Acetylcholine alters the amplitude and frequency of gut motility. (A) Gut motility frequencies for 6 dpf control larvae (blue, n=31) and larvae immersed in 2.5 mg/ml acetylcholine (ACh; orange, n=30), showing an increased frequency in ACh-treated larvae (mean ± s.e.m. = 2.38 ± 0.0331 min^-1^) compared to untreated controls (2.23 ± 0.0527 min^-1^). Each point represents data from a five-minute video of a single larva, captured at five frames per second. Darker circles and lighter diamonds represent two independent experiments. (B) Gut motility amplitudes corresponding to the same experiments depicted in panel (A). Both the mean and the standard error of the mean of gut motility amplitudes for ACh-treated larvae (0.128 ± 0.0174 μm) are higher than controls (0.0952 ± 0.0121 μm).

To further examine the utility of our program, we extracted information about the wave propagation speed and amplitude of motility events. ACh-treated larvae exhibited no difference in wave speed compared to controls (Suppl. Fig. 1). However, 2.5 mg/ml ACh increased the median motility amplitude by over 50% (Fig. 2B). The mean ± s.e.m. amplitude values at 0.2 seconds per frame were 0.128 ± 0.0174 μm and 0.0952 ± 0.0121 μm for ACh-treated and control larvae, respectively. Increased larval gut contraction strength has not been reported previously, but is reminiscent of similar ACh-induced effects seen in *ex vivo* smooth muscle preparations from adult zebrafish (Holmberg et al., 2004).

### Feeding increases gut motility frequency and sustains amplitude during development

Food is well known to influence gut motility, in particular by triggering contractile waves often referred to as peristaltic motions (Furness, 2006; Olsson and Holmgren, 2011). In zebrafish, the influence of food on gut motility patterns has not previously been assessed. We predicted that food-induced contractions would lead to observable and quantifiable increases in motility amplitude. To test this hypothesis, we compared gut motility parameters in both fed and unfed siblings over three days of development from 5-7 dpf. As before, 5 minute DICM videos were taken at 5 fps and analyzed. Videos in which food pieces were evident within the gut were discarded, as velocimetry is unable to distinguish cellular movement from food movement.

We compared gut motility frequency (Fig. 3A) and amplitude (Fig. 3B) in both fed and unfed siblings. Surprisingly, we found that feeding larvae alters the frequency of gut motility. At 5 dpf, there is little difference between fed and unfed larvae (Fig. 3A). However, for the next two days of integrated food consumption, fed gut motility frequency diverges away from that of unfed siblings (2.24 ± 0.03 min^-1^ and 2.06 ± 0.03 min^-1^ for 6 dpf fed and unfed, respectively, and 2.45 ± 0.06 min^-1^ and 2.10 ± 0.06 min^-1^ for 7 dpf). Strikingly, whereas unfed larvae appear to have monotonically decreasing frequency with age, fed larvae show higher gut motility frequency at 7 dpf than at 6 dpf (Fig. 3A).

**Figure 3:**
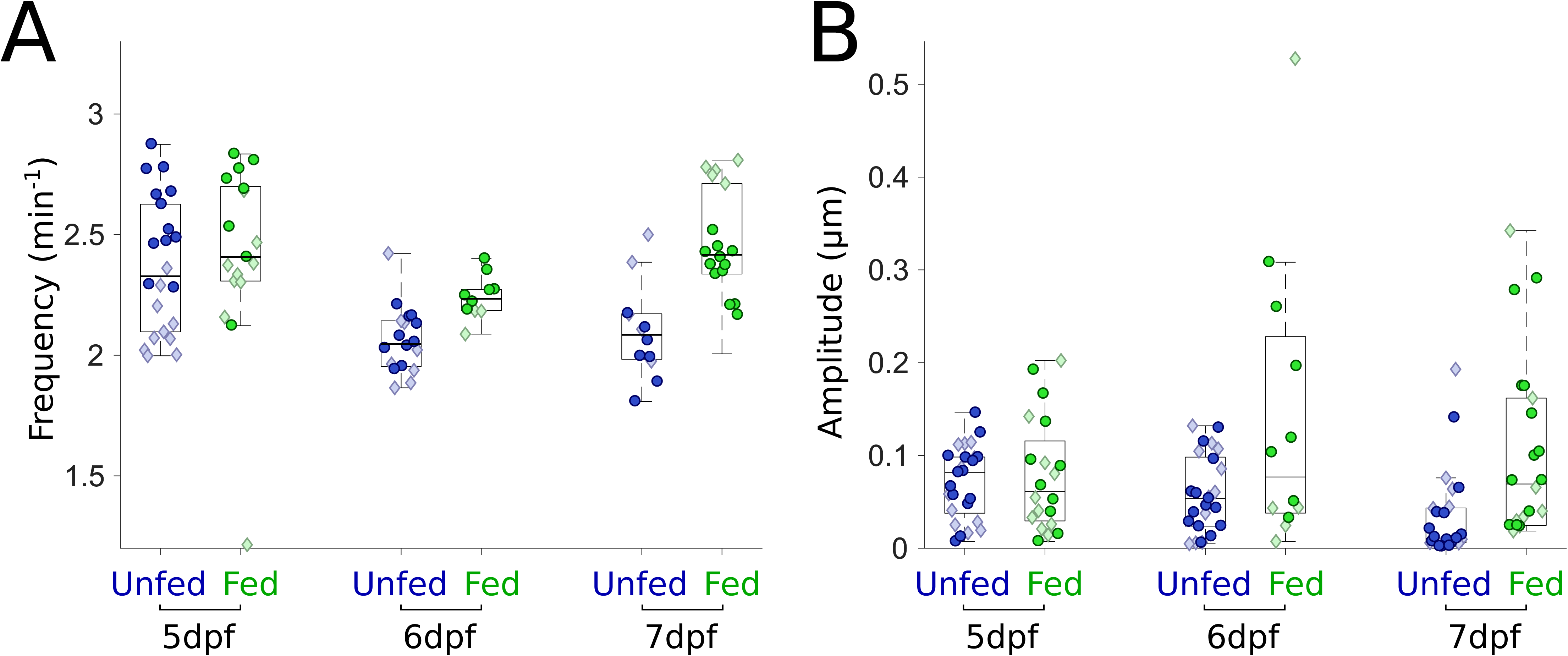
Feeding increases the frequency and amplitude of gut motility. (A) Gut motility frequencies for unfed (blue, n=22,18,12) and fed (green, n=17,10,18) larvae over three days of development. Frequencies of fed and unfed larvae remain similar after one day of feeding. Frequencies become different over the next two days, with fed larvae showing higher frequencies. Darker circles and lighter diamonds represent two independent experiments. (B) Gut motility amplitudes corresponding to the same experiments depicted in panel (A) for unfed (blue, n=25, 25, 25) and fed (green, n=20,12, 22). As in panel (A), amplitudes are similar to one another one day after feeding but the means become significantly different over the next two days.

As expected, zebrafish gut motility amplitude increased with feeding, though in an age-dependent manner (Fig. 3B). At 5 dpf, after one day of feeding, little change in amplitude is evident (Fig. 3B). However, for the next two days of integrated food consumption, the amplitude difference between fed and unfed larvae increases, with the median value in fed larvae being 1.4 times greater than in unfed larvae at 6 dpf, and 6.5 times greater at 7 dpf. The mean ± s.e.m. values were 0.143 ± 0.045 μm and 0.058 ± 0.008 μm for fed and unfed larvae respectively at 6 dpf, and 0.103 ± 0.021 μm and 0.033 ± 0.009 μm at 7 dpf. At both 6 and 7 dpf, feeding also leads to an increased spread in the amplitude data (Fig. 3B).

### Larvae lacking ENS innervation display decreased motility amplitude

Changes in ENS innervation are known to affect gut motility (Heanue et al., 2016; Kuhlman and Eisen, 2007; Uyttebroek et al., 2016). We analyzed gut motility parameters in 5-7 dpf *ret^hu2846/hu2846^* (hereafter referred to as *ret^-/-^*) mutants. These fish lack ENS innervation and serve as models for Hirschsprung disease, a human congenital ENS disorder. Surprisingly, we found no discernible difference in frequency (Fig. 4A) or wave velocity (Suppl. Fig. 2), in contrast to a recent study reporting reductions in these parameters in 7 dpf *ret* mutant larvae (Heanue et al., 2016). However, we found that on average, zebrafish *ret* mutants show reduced gut motility amplitudes compared to wild-type siblings at all days examined (Fig. 4B). We previously noted the lower motility amplitude of *ret* mutants, using an early version of this analysis approach (Wiles et al., 2016).

**Figure 4:**
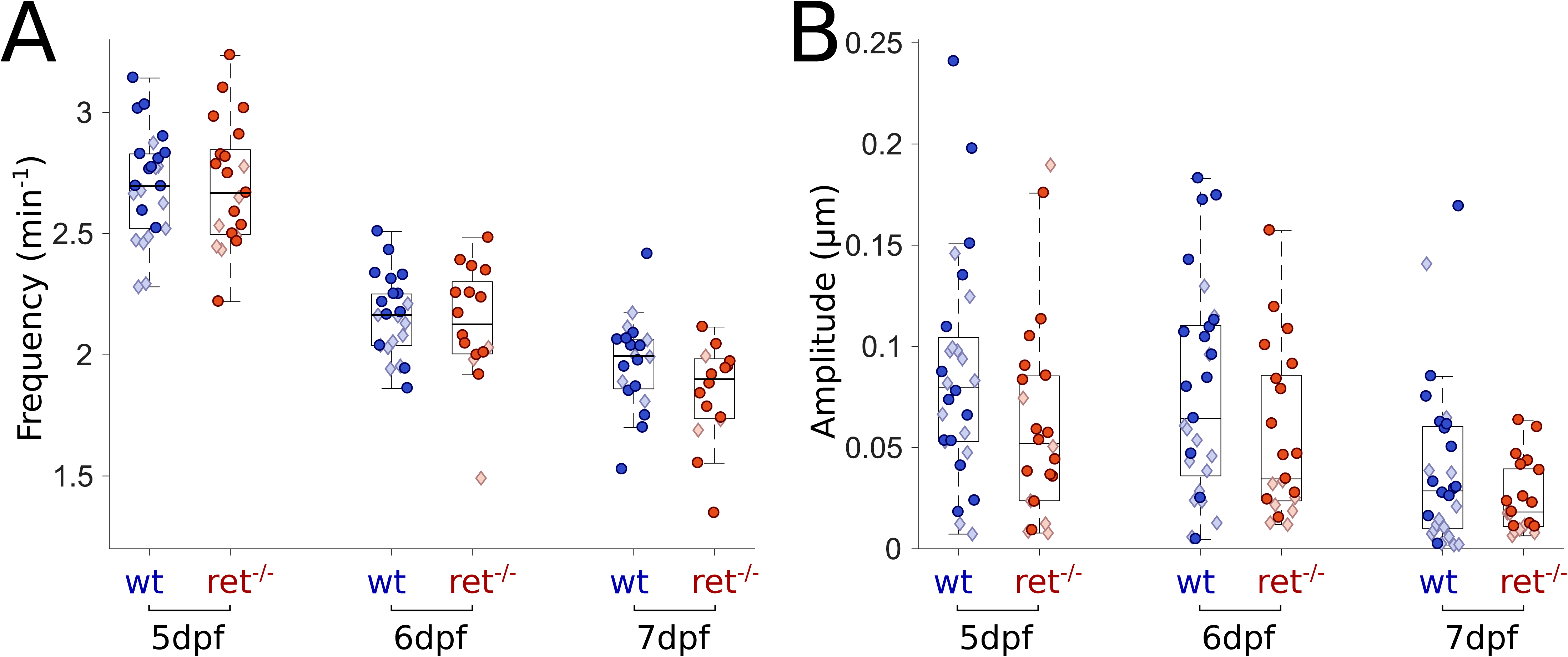
*ret* mutants lacking an ENS display similar frequencies and reduced amplitudes compared to wild-type siblings. (A) Gut motility frequencies for wild-type (wt) (blue, n=25, 23, 20) and *ret^-/-^* (red, n=21,16,16) larvae over three days of development. Frequencies of *ret^-/-^* and wt siblings are the same over three days of development. Darker circles and lighter diamonds represent two independent experiments. (B) Gut motility amplitudes corresponding to the same experiments depicted in panel (A) for wt (blue, n=28, 29, 28) and *ret^-/-^* (red, n=22, 21, 21). Amplitudes and standard deviations of those amplitudes of *ret* mutants are consistently lower over all three days compared to wt.

### Variability in gut motility parameters is dependent on the ENS

The amplitudes of gut motility events show considerable variability between individuals, especially among wild-type larvae (Fig. 4). We hypothesized that this variability would also be manifested within individuals over longer observation times, and that it would be larger in wild-types than in *ret* mutant larvae. To test this hypothesis, we imaged 6 dpf larvae for approximately 90 minutes, and analyzed the resulting gut motility patterns as described above, generating spectral signatures of 4 minute sliding windows spanning the full duration of the movies (Fig. 5A,B). We found that wild-type larvae show a remarkable range of amplitudes over time both within and between individuals (Fig. 5A,C). In comparison, *ret* mutant larvae display much less amplitude variability within individuals (Fig. 5B,D).

**Figure 5:**
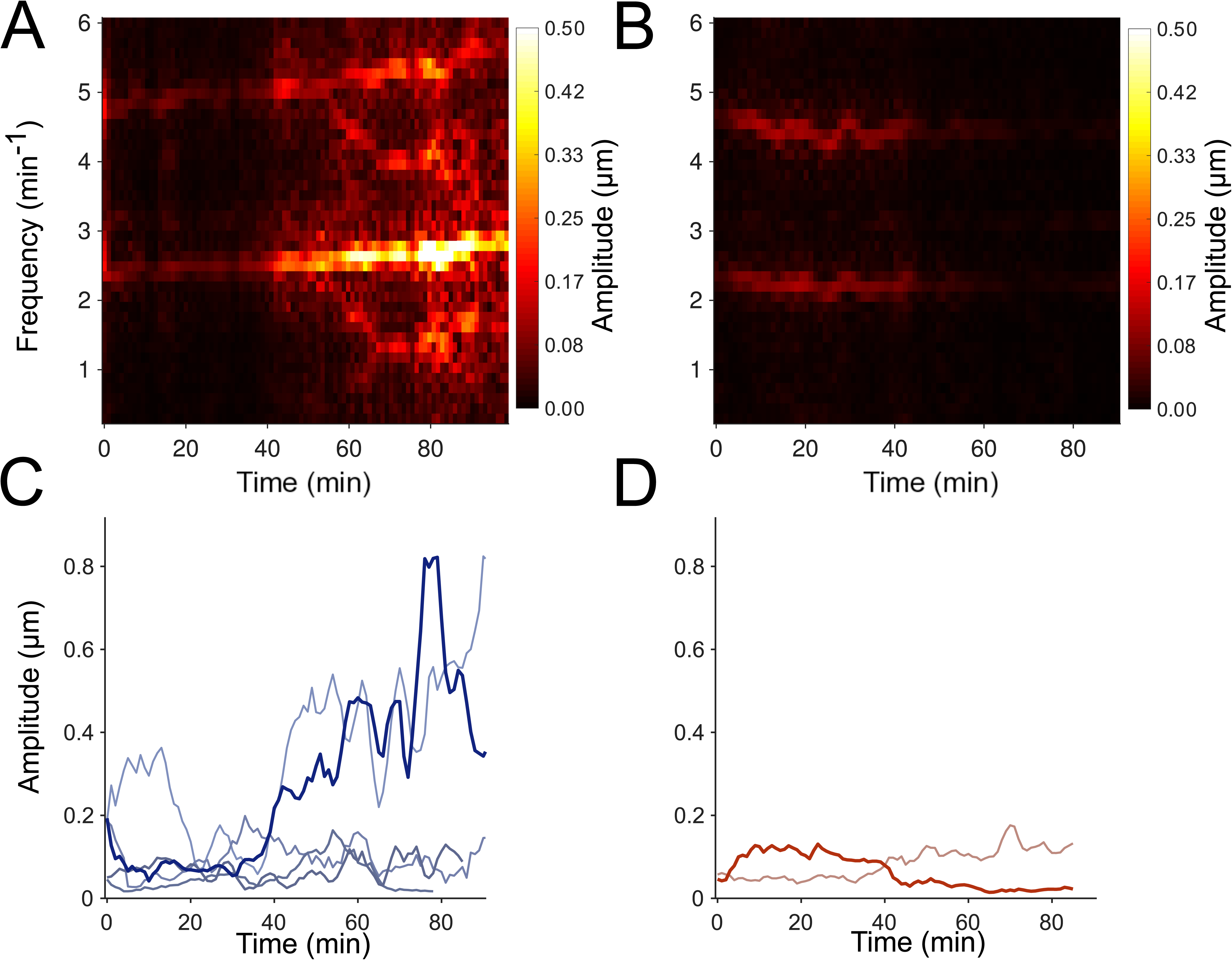
Wild-type larvae have higher amplitude variability than *ret* mutants. (A) Spectrogram illustrating the time-varying gut motility power spectrum of a wild-type (wt) larva over 1.5 hours. Each column depicts the power spectrum calculated over a 4-minute window. (B) A spectrogram of a single *ret* mutant larva. (C) Maximum Intensity Projections (MIP) of spectrograms for wt larvae (n=5); each curve represents a different larva. The bolded blue curve is the MIP of the spectrogram provided in (A). (D) MIP of the spectrograms for *ret* mutant larvae (n=2). The bolded red curve is the MIP of the spectrogram provided in (B). The amplitudes for both *ret* mutant larvae are lower and less variable over time than most wt larvae.

## Discussion

The complex motility patterns of the vertebrate gut are crucial to its function, and are modulated by developmental processes, physical and chemical stimuli, and the pathology involved in a variety of disease states. The question of how to measure and characterize gut motility in a way that captures its essential features is therefore both important and timely.

Periodic, propagative contractions are critical for gut activity. For any periodic oscillatory motion, frequency and amplitude are essential and distinct characteristics. It has long been realized that data from imaging studies can readily yield gut motility frequencies and related properties such as wave propagation speeds, for example via image-derived STMaps. Straightforward yet robust amplitude measures have proven more challenging to obtain. We therefore developed and assessed a new image analysis approach that combines image velocimetry, commonplace in studies of fluid dynamics, and spectral analysis, ubiquitous in signal processing applications, to provide quantitative measures of parameters related to both frequency and amplitude.

We apply this approach to data derived from DICM imaging of the larval zebrafish gut. DICM is well-suited to this analysis as it provides high contrast images of sub-cellular features as well as intrinsic optical sectioning. The former facilitates image velocimetry, as there are abundant features to correlate between video frames, while the latter avoids blurring and averaging over the depth of the sample.

We assessed our method in known as well as novel settings including ACh treatment, comparing fed to unfed zebrafish larvae, and analyzing zebrafish mutants lacking ENS innervation. Previous studies in zebrafish using conventional STMaps have found differences in parameters such as gut peristaltic frequency and the speed at which peristaltic waves travel along the gut for various phenotypes (Heanue et al., 2016; Rich et al., 2013; Uyttebroek et al., 2016) and experimental conditions (Holmberg et al., 2007; Holmberg et al., 2006; Holmberg et al., 2004). We have shown, however, that there exist phenotypes that are identical in frequency or wave speed that are nonetheless different in the amplitude of gut motility, highlighting the importance of examining this axis of behavior. In addition, even for known experimental settings like treatment with ACh, we found that in addition to the expected increase in frequency, the amplitude of gut movements is also increased. Our program allows a more comprehensive analysis of gut motility parameters. Furthermore, the framework of cross-correlations, spectral analysis, and open-source software enables additional parameter extractions, if desired. The image analysis method presented here quantifies imaged motions in an automated and reproducible manner. In addition, it is agnostic to the types of images it analyzes, making it versatile for a variety of cellular movements.

Due to the indiscriminate and automated nature of the analysis, a wider range of movements will be recorded when compared with methods that make use of manual feature identification. Consequently, some of the parameters defined in this study may not correspond directly to parameters obtained in previous research. As an example, previous studies have defined the frequency of gut motility only when a sustained wave travels along a large enough distance of the gut. In contrast, our method will identify the frequency of any periodic motion, whether it is a standard motility event or a single muscle cell firing repeatedly. In future applications, the user could define their own parameters from the QSTMap or from even the raw velocity vector field.

Our observation of increased gut motility frequency and amplitude in fed, compared to unfed, larval zebrafish provides the first assessment of how feeding alters motility in these animals. It is well-known in general that specific gut movements are triggered by food (Furness, 2006; Olsson and Holmgren, 2011). In mammals, the gut either displays stationary contractions that are non-propulsive and are necessary for mixing food or propulsive contractions that transport gut contents (Furness, 2006; Wood, 2008). Our findings point to rich dynamics that can be rigorously studied in zebrafish, varying for example the duration and type of feeding. Food may also shape the microbial composition of the gut, as recent work has shown that apparent inter-microbial competition can be governed by gut motility (Wiles et al., 2016) and that zebrafish mutants with altered motility assemble communities that can be distinguished by abundance of particular members (Rolig et al., 2017).

Our examination of gut motility parameters in *ret* mutant zebrafish larvae lacking ENS innervation highlights both the utility of our analysis and the complexity of mechanisms underlying gut motility. Heanue and colleagues (2016) examined gut motility parameters in 7 dpf *ret* mutant larvae and found reduced frequency, contraction distance and contractile velocity compared to wild-type siblings (Heanue et al., 2016). In contrast, our study found a noticeable difference in amplitude but did not find differences in frequency or speed between 5 dpf and 7 dpf. The mutant allele used in these two studies is the same (*ret^hu2846^*) and the mutant larvae show the same phenotype regarding enteric neurons, namely a total lack of neurons except for a few in the intestinal bulb [Suppl. Fig 3, (Heanue et al., 2016)]. However, the two mutant lines have been maintained on different wild-type backgrounds [Tubingen Longfin, (Heanue et al., 2016), AB (our study)]. Additionally, in contrast to the study of Heanue and colleagues (2016), we do not observe an ENS phenotype with fewer neurons and altered gut motility in heterozygous larvae. One possible explanation for the difference in the gut motility defect is differences in the genetic background due to differences in the wild-type lines, most likely related to the high degree of heterogeneity in the number of SNPs between different genetic backgrounds (LaFave et al., 2014). Interestingly, this difference is very reminiscent of Hirschsprung disease, a genetically complex disorder that displays significant phenotypic variation, for example differences in the extent of intestinal aganglionosis, even among individuals with the same mutant alleles (Heanue et al., 2007). These results highlight the importance of detecting complementary and independent gut motility parameters, as in our *ret* mutants on the AB background only amplitude was affected. If we had analyzed our data using established methods, we would have concluded that gut motility parameters did not differ between in *ret* mutant larvae and their wild-type siblings. Thus, this newly developed approach provides additional parameters that may be differentially affected in different enteric neuropathies or gut diseases.

Phenotypic variation is a hallmark of ENS diseases such as Hirschsprung disease. We observe in general a striking degree of variability within and among individual zebrafish larvae with regard to gut motility amplitude (Fig. 5). This variability is displayed in all the data measurements, but becomes most apparent during longitudinal imaging. It has been previously reported that the speed of gut transit varies considerably among individuals (Field et al., 2009). We suggest that the variability represents different gut motility modes that reflect different gut states at any given time point, for example the difference between when food is being mixed and when nutrients are being absorbed, which is then reflected in amplitude differences. Whereas two of the larvae shown in Fig. 5 show strong, varying increases in amplitude, the three other larvae show moderate changes in amplitude. In contrast, *ret* mutant larvae show very little change in amplitude over time. The ENS provides the intrinsic gut innervation that regulates gut movements (Furness, 2006). We propose that these amplitude changes are regulated by the ENS and may thus be absent from *ret* mutant larvae, motivating future work to establish connections between gut motility modes and specific ENS neuronal activity, and their alteration in the course of gut diseases.

## Materials and Methods

### Zebrafish Husbandry

All experiments were carried out in accordance with animal welfare laws, guidelines and policies and were approved by the University of Oregon Institutional Animal Care and Use Committee. Wild-type and *ret^hu2846^* embryos were allowed to develop at 28.5°C and staged by hours post fertilization according to morphological criteria (Kimmel et al., 1995). Wild-types and *ret^hu2846^* were of the AB background.

### Imaging Experiments

Specimen mounting was performed as described previously (Jemielita et al., 2014). Briefly, larvae were anesthetized in 80 μg/ml tricaine methanesulfonate (Western Chemical, Ferndale, WA) for several minutes at 28°C. Larvae were then immersed in a liquified 0.5% agar gel (maximum temperature 42°C) and drawn into a glass capillary. The gel, once solidified, was mounted onto a microscope imaging chamber containing embryo medium (EM) with 80 μg/ml tricaine methanesulfonate maintained at 28°C. The solidified gel and the larva were extruded into the imaging path to prevent the capillary glass from interfering with imaging. The mid-region of the gut was imaged, approximately 200 μm anterior of the anus (vent).

Imaging was performed using a custom-designed and custom-built microscope capable of differential interference contrast microscopy as well as light sheet fluorescence microscopy (Baker et al., 2015). The specimen was illuminated by a polarized 447nm LED (Quadica Developments Luxeon Star Brantford, Ontario, Canada) and imaged using a standard microscope objective (Zeiss Oberkochen, Germany DICMMPlan Apochromat, 40x/1.0). A Nomarski prism and polarizer were oriented in such a way as to provide Differential Interference Contrast (DIC) (Baker et al., 2015). The resulting image was then focused onto a sCMOS Camera (Cooke, Kelheim, Germany, pco.edge). Movies were taken with 1 ms exposure times at 5 frames per second.

### Acetylcholine treatment

Wild-type larvae were raised in EM until 6 dpf. Acetylcholine treatments were essentially performed as previously described (Shi et al., 2014). Briefly, larvae were individually transferred to EM containing either 0.5% DMSO or 0.5% DMSO with Acetylcholine Chloride (Sigma-Aldrich, A6625; 2.5 mg/mL). Larvae were exposed to these conditions for a total of 20-30 minutes, anesthetized with tricaine for several minutes at 28°C, and mounted for imaging as described above.

### Feeding of zebrafish larvae

Wild-type larvae were raised in EM until 4 dpf and transferred to embryo medium at 5 parts per thousand salinity (E5) in a new dish and rotifers added to the dish. Fresh rotifers were added at 5 dpf and 6 dpf, so the fed zebrafish larvae had food *ad libidum*; 7 dpf fish were provided food for three days. Larvae were examined to ensure they had no food in their gut immediately prior to imaging, as PIV may track gut contents, such as food, instead of the gut wall.

### Particle Image Velocimetry (PIV) and Quantitative Spatiotemporal Maps (QSTMaps)

PIV is a well-established image analysis technique that takes as its input a set of images and outputs a corresponding set of velocity vector fields representative of the motion contained within those images. To perform PIV, we used publicly available software called “PIVLab” [http://pivlab.blogspot.com] in addition to several home-built Matlab programs, provided as Supplementary Material.

A comprehensive description of how PIV works, its many different implementations, and how it is optimized can be found elsewhere (Willert and Gharib, 1991). However, a simple example, representative of the key features of the technique, is as follows: A two-dimensional image *I*_p_(*x*,*y*) (known as an “interrogation area,” possibly the subset of an even larger image) at frame *p* of an image series is subdivided into a grid. We denote the subset of *I*_p_(*x*,*y*) centered at grid element (i,j) as the template *t*_p,ij_(*x*,*y*). For each template in frame *p*, the cross correlation with the frame (p+1) is calculated:

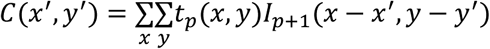

The location of the maximum of *C*(*x*’,y’) gives the most likely displacement of that template neighborhood from one frame to the next. The x’ and y’ at which C is maximum for each template correspond to each of the red arrows in Fig. 1B and Suppl. Movie 1. This is repeated over all grid elements to generate a displacement or velocity vector field, and then is repeated over all pairs of frames.

For this study, we used a first pass template size of 32 pixels corresponding to 20.8 microns in the image plane. Preliminary preprocessing consists of using PIVLab’s built in PIVlab_preproc function, using contrast enhancement (CLAHE, size 50) and a high pass filter (size of 15 pixels). PIVLab then performs PIV over the entire image, segregating the resultant velocity vector field into a grid whose vertices are separated by 32 pixels. After this processing, a user defined mask is applied to the region of interest (in our case, an area containing the gut) and vertices outside of the mask are discarded.

As the geometry of the gut is not conserved in space or between individual larvae, masking results in the remaining vertex positions and numbers being spatially inconsistent from one data set to another and difficult to deal with numerically. To manage this, a new grid is generated to better accommodate the unique geometry. This new grid has a constant number of rows and columns and is distributed inside the mask in such a way as to fill most of the area. To do this, a curve is drawn by the user which represents the centerline of the mask (not necessarily the geometric center; in our case, the gut lumen). At each discrete position along the curve (equal in distance to the original PIV spacing), a constant number of vertices is distributed orthogonal to the curve at that position. This results in axes which, while spatially varying, are a better representation of the DV and AP axes of the gut (Suppl. Movie 1). The original velocity field is transformed to the new grid by bilinear interpolation and is its components are projected into the local DV and AP components.

We generate a QSTMap from the resulting velocity field. We are primarily concerned with the AP component of motion, and its variation along the AP axis. We therefore average the AP component of the frame-to-frame displacements along the DV direction, resulting in a one-dimensional map of displacement as a function of AP position, for each point in time (represented in the bottom half of Suppl. Movie 1). Plotting these functions over time gives the QSTMap. A representative data set is shown in Figure 1C. We note that other analyses are possible, for example considering DV displacements, which can be implemented by modifying our code.

### Cross-Correlation Plots Define Frequency and Wave Speed

Larval gut motility waveforms can be individualized and complex. For most cases, the velocity waveform does not have a well-defined set of maxima that can clearly be followed across position and time. These waves, however, often have similar structures that repeat over time. Because of this, we take the QSTMap, *Q*(*x*,*t*), and apply the cross correlation

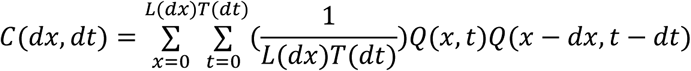

where *L*(*dx*) = *L_0_* - dx is the length of the gut that can be examined for a given offset *dx*, *L_0_* is the total AP length of the analyzed gut segment (typically around 400 μm), T(dt) = T_0_ - dt is the time that can be examined for a given offset dt, T_0_ is the total time of the analyzed video (typically around 5 minutes).

An example of the resulting cross correlation is shown in Figure 1D. Even for multi-modal waveforms, the cross correlation results in a well-defined set of maxima that linearly increase over changes in distance. Therefore, we find the locations of the maximum of *C* and fit it to a line. The inverse slope of this line (Figure 1D, orange arrow) is defined as the wave speed.

The first non-zero peak in the autocorrelation of a signal is the time at which velocities at any position in the gut are most similar with themselves. The location of this peak therefore provides a robust measure of the frequency (Figure 1D, green bracket).

### Spectral Analysis Defines Amplitude

To define amplitude, we needed a measure that is robust against noise and that focuses on the periodic peristaltic events and ignores occasional large vectors that result from motions such as larvae moving. We therefore perform a Fourier Transform of our QSTMap at each AP position, transforming each *Q*_x_(*t*) into a function of frequency, *f*, rather than time:

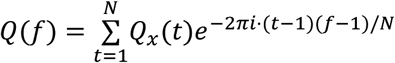

To obtain the signal strength (“power”) at any frequency, we take the modulus (the signal multiplied by its complex conjugate) of Q_x_(*f*). Having previously found the frequency of gut motility from the cross correlation, we define the amplitude as the square root of the power at the frequency of gut motility. For simplicity, our analysis considers only the average of these values over the entire gut in the field of view. Figure 1E, shows the resultant power spectrum of the QSTMap from Figure 1C, with the red box outlining the peak power at the frequency of gut motility.

## Acknowledgements

The authors thank Brandon Schlomann, Savannah Logan, Eric Corwin, and Karen Guillemin for useful discussions. Research reported in this publication was supported by the NIH as follows: NIGMS award P50GM098911 and NICHD award P01HD22486. The content is solely the responsibility of the authors and does not represent the official views of the NIH. Research reported in this publication was also supported by the National Science Foundation under award numbers 0922951 and 1427957, the M.J. Murdock Charitable Trust, and a REACHirschsprungs Foundation Research Grant (to JG).

## Figure legends

**Supplementary Figure 1: Acetylcholine does not alter the wave speed of zebrafish gut motility.** Wave propagation speeds for 6 dpf control larvae (blue, n=31) and larvae immersed in acetylcholine [ACh; 2.5mg/ml (orange, n=30)]. As larger velocities tend to be unreliable due to the low temporal resolution of our data, measured velocities are capped at a user-defined threshold, given by the dashed line. Each point is derived from a five minute video of a single larva. Darker circles and lighter diamonds represent two independent experiments.

**Supplementary Figure 2: *ret* mutant zebrafish larvae show no noticeable difference in gut motility wave speed compared to wild-type (wt) siblings.** Wave propagation speeds for wt (blue, n=25, 23, 20) and *ret^-/-^* (red, n=21, 16, 16) larvae over three days of development. As larger velocities tend to be unreliable due to the low temporal resolution of our data, measured velocities are capped at a user-defined threshold, given by the dashed line. Each point is derived from a five minute video of a single fish. Darker circles and lighter diamonds represent two independent experiments.

**Supplementary Figure 3: *ret* mutant larvae lack ENS innervation.** Lateral views of 6 dpf sibling larvae, from combined brightfield and fluorescence images. (A) Wild-type larva with ENS neurons expressing GFP driven by the *phox2b* promoter (*phox2b*:GFP) along the entire length of the gut. (B) *ret* mutant larva, which lacks ENS innervation except for a few GFP-positive ENS neurons in the anterior-most part of the gut (arrows). Scale bar = 100μm in A, B.

**Supplementary Movie 1: Larval gut motility and Particle Image Velocimetry.** (Top) DIC movie of larval zebrafish gut motility with PIV vectors overlaid in red. The magnitude of the vector represents the instantaneous velocity of a small section of the gut and the angle represents the direction it is traveling. Total time: 22 seconds. (Bottom) Averaging the anterior posterior component of the velocity along the dorsal-ventral direction generates a single curve at each time point. QSTMaps are the surfaces generated by these curves over time.

